# Characterization of a new laboratory colony of *Anopheles funestus* mosquitoes established in Ifakara, Tanzania

**DOI:** 10.1101/2025.11.23.690074

**Authors:** Emmanuel E. Hape, Rukiyah M. Njalambaha, Letus L. Muyaga, Ismail H. Nambunga, Joseph P. Mgando, Dickson D. Mwasheshi, Neema K. Nombo, Daniel M. Mabula, Munyaradzi P. Zengenene, Najat F. Kahamba, Joel O. Odero, Halfan S. Ngowo, Salum A. Mapua, Prosper P. Chaki, Nicodem J. Govella, Issa N. Lyimo, Samson S. Kiware, Dickson W. Lwetoijera, Brian B. Tarimo, Emmanuel W. Kaindoa, Prashanth Selvaraj, Frederic Tripet, Charles S. Wondji, Francesco Baldini, Lizette L. Koekemoer, Heather M. Ferguson, Fredros O. Okumu

## Abstract

**Background:** *Anopheles funestus*, a major vector of malaria in Africa, has proven difficult to colonize in laboratory settings, impeding research on its biology and control. After several attempts, our team recently succeeded in colonizing a strain of *An. funestus* from Tanzania (FUTAZ). The objective of this study was to analyse the key fitness and genotypic characteristics of these mosquitoes during multiple filial generations of laboratory adaptation and compare them to wild *An. funestus* from Tanzania and a pre-existing colony of *An. funestus* from Mozambique (FUMOZ).

**Methods:** Measures of mating success (percentage of female mosquitoes inseminated), body size (wing length), fecundity (number of eggs laid per female), and insecticide susceptibility (percentage of 24-hour mortality after exposure to insecticides) were compared between the newly established colonies of Tanzanian *An. funestus* (FUTAZ colonies), the long-established FUMOZ colonies, and a colony of *Anopheles arabiensis* maintained in the same laboratory. The maternal lineages of the *An. funestus* mosquitoes were investigated through a hydrolysis probe analysis of their mitochondrial DNA to identify distinct clades, I and II. Additionally, other intragenomic variations were examined through a PCR analysis of restriction fragment length polymorphisms (RFLP) on the third domain of 28S ribosomal DNA. These molecular markers were used to compare the FUTAZ colonies, FUMOZ colonies in Tanzania and South Africa, and the wild-collected *An. funestu*s from Tanzania.

**Result:** The mating success and body size of FUTAZ females declined significantly from filial generations F1 to F6 relative to the founder population (F0), but then increased from F7 onwards eventually matching FUMOZ by F9. Fecundity was similar across all colonies tested. However, it took significantly longer for 50% of the females in the FUTAZ and FUMOZ colonies (over 10 days) to mate compared to females in the *An. arabiensis* colony (approximately 5 days). Insecticide resistance appeared to be lost during colonization, but this varied with insecticide classes. Majority of mosquitoes in the FUTAZ colony, as well as the wild-caught Tanzanian *An. funestus* belonged to Clade I (80.4-89.4%) and RFLP type “Y” (90.5-91.4%), while the FUMOZ colonies were mostly Clade II (65.5-88.5%) and RFLP type “MW” (90.5-91.5%).

**Conclusion:** This study suggests that the mating success and body size of *An. funestus* decreases significantly during the early stages of colonization, then increase as the mosquitoes adapt to laboratory conditions. It is therefore crucial to have a large enough founder population to persist through these early generations in order to achieve stable colonization of *An. funestus*. The Clade and RFLP genotyping demonstrated the genetic similarities between the FUTAZ mosquitoes and wild-caught Tanzanian *An. funestus*, but also showed that the new colony can be distinguished from the FUMOZ colony.

## Background

The burden of malaria has significantly decreased since 2000 [1], primarily because of key interventions such as insecticide treated nets (ITNs), indoor residual spraying (IRS) and improved case management [2,3]. However, there were still over 247 million cases and 619,000 deaths due to the disease in 2021 [4], suggesting significant improvements are necessary to achieve elimination.

In sub-Saharan Africa, malaria is primarily transmitted by three members of the *Anopheles gambiae sensu lato* species complex (*An. gambiae sensu stricto*, *An. coluzzii* and *An. arabiensis*), and *An. funestus s.s.* (hereafter referred to simply as *An. funestus*) [5]. Rearing and maintaining these mosquitoes under laboratory conditions (colonies) is important to facilitate research on their biology and control. Since mosquitoes in the *An. gambiae s.l.* are relatively easy to colonize, much has been learned about their ecology and response to interventions from laboratory studies. In contrast, comparatively little is known about *An. funestus* because of the inability to reliably and repeatedly colonize this species under laboratories conditions [6–9]. Existing protocols for colonization of *An. funestus* have been broadly unsuccessful in establishing new colonies from the wild [10–12]. Despite numerous attempts, only two lines of *An. funestus* have been successfully colonized from the wild; one from Mozambique (FUMOZ, in 2000) and another from Angola (FANG, in 2002) [13]. There are also reports of an earlier successful colonization of this species by Service and Oguamah in Nigeria in 1950s, but this colony no longer exists [14].

Detailed analysis of mosquito fitness traits during laboratory rearing can provide valuable insights into the main barriers to colonization, and time required for adaptation in the controlled environments [15]. Already, multiple bottlenecks have been identified as possibly hindering the laboratory colonization of *An. funestus*, many of which are associated with important life cycle processes such as egg-laying, larval development, and adult survival [10–12]. Obstacles in colonizing *An. funestus* are also partly due to eurygamy, a common feature of many *Anopheles* species [16], defined as the inability to mate in confined laboratory settings. Forced artificial mating has been reported as a successful technique for establishing colonies of eurygamous anophelines [16,17]; however, it is time demanding and requires highly-skilled personnel [18]. Similarly, a forced oviposition technique has been used, where individual gravid mosquitoes are kept in small tubes or oviposition cups lined with wet filter papers to enhance egg laying [9,19,20]. The forced oviposition method has been particularly useful for producing offspring from wild-collected *An. funestus* females, but is insufficient on its own for establishing stable colonies over multiple generations [9,11,21]. Previous studies have used the forced oviposition approach to characterize fitness differences between wild females and their offspring when reared in the laboratory for *An. funestus* [11], and other *Anopheles* [22]. These studies provide insights into how *Anopheles* species adapt to laboratory conditions over multiple generations [22,23].

Several mosquito genetic and phenotypic traits have been observed to change within the first few generations of colonization. For example, mosquito lines become highly inbred within a few generations of colonization [24,25]. Additionally, traits that confer a fitness advantage in the laboratory (high mating success and survival in cages) are expected to be selected during colonization. In contrast, some mosquito traits that provide a selective advantage in the field but not in the laboratory may be lost during colonization [26]. For example, the extensive use of pesticides in both agriculture and public health in Africa has generated strong insecticide resistance in many malaria vector species including *An. funestus* [27,28]. While these resistance traits are crucial for survival in the wild, they may be costly to maintain in the absence of continued selection [29,30]. Consequently, resistance traits are likely to be lost in laboratory colonies where selection for resistance is removed [31,32].

Besides phenotypic attributes, it is also vital to conduct detailed genetic characterization of new mosquito colonies to understand their closeness and differences from either the wild populations or existing colonies. Various methods have been used to distinguish between members of the *An. funestus* group. These include variations on the third variable domain (D3) of the nuclear 28S ribosomal-DNA (rDNA) [33,34], and analysis of the internal transcribed spacer 2 region (ITS2) of the rDNA, which is the basis for current species differentiation [33, 34]. Six sibling species in the *An. funestus* group, including *An. rivulorum*, *An. rivulorum-like*, *An. parensis*, *An. funestus*, *An. vaneedeni* and *An. leesoni*, can be identified using a species-specific PCR established by Koekemoer *et al*., [36] and Cohuet *et al*., [35]. Moreover, genetic structuring in the populations of *An. funestus* has been observed, and two cryptic subdivisions described based on mitochondrial DNA clade analysis targeting the NADH Dehydrogenase subunit 5 (ND5) and Cytochrome Oxidase I (COI) [35, 36]. These assays have revealed that *An. funestus* populations subdivide into Clade I, which is widespread across Africa and Clade II, previously known to be only in Mozambique, Madagascar, and Zambia [37–39]. Clades I and II are reported to have evolved independently for ∼1 million years [38]. In addition, microsatellite DNA markers have shown major genetic heterogeneities between the populations of *An. funestus* from coastal and western Kenya [34], and low gene flow between *An. funestus* populations from eastern or southern African, and those from Malawi [34]. Lastly, a PCR- restriction fragment length polymorphism (PCR-RFLP) has also been developed to distinguish between *An. funestus* populations based on geography, and to understand the effects of physical barriers such as the Great Rift Valley on population-level gene flow [33,34]. Such analysis has previously identified geographically segregated *An. funestus s.s*. populations belonging to either M, W, MW, Y or Z- types based on the PCR-RFLP profiles [33,34], supporting distinct population structures between regions.

Until recently, studies of *An. funestus* fitness during colonization were not feasible given the difficulties of laboratory adaptation. No measurements of mosquito fitness were taken during the early stages of colonization of the only two stable *An. funestus* lines (FUMOZ and FANG), and a recent attempt to measure the fitness of a Tanzanian *An. funestus* line was limited as that colony persisted for only two generations in the laboratory [11]. However, after several attempts, our team recently succeeded in colonizing a strain of *An. funestus* from Tanzania (FUTAZ). This colony was established by first building up large ‘F1 populations’ in the laboratory by repeatedly adding eggs from wild female founder populations (F0) collected over 26 months from two sites in Tanzania (May 2018 to July 2020). Once a sufficiently large number of individuals had been built up in the F1 cages, a stable laboratory population was established which has now been maintained in the laboratory for over 30 generations without addition of further wild offspring.

The aim of this study was therefore to characterize the major fitness traits of the recently established *An. funestus* colony at Ifakara Health Institute during multiple generations of laboratory adaptation. Additionally, we used the proven molecular markers for Clade [37] and PCR-RFLP [34] analysis to examine genetic similarities and differences between the FUTAZ colony, wild-collected *An. funestus*, also from Tanzania and the pre-existing colony of *An. funestus* originally from Mozambique (FUMOZ).

## Material and Methods

### Biological material

The experiments described here were conducted using: a) field-collected *An. funestus* from two rural Tanzanian villages (Wild FUTAZ, Ikwambi (− 7.97927° S, 36.81630° E) and Tulizamoyo (− 8.35447° S, 36.70546° E)), b) laboratory colony established using the Wild FUTAZ, c) laboratory colony of *An. funestus* mosquitoes originally colonized in 2000 using field-collected samples from Mozambique (FUMOZ) and maintained by the Vector Control Reference Laboratory (VCRL) and National Institute of Communicable Diseases (NICD) in South Africa [13], and d) a FUMOZ colony maintained in Tanzania.

Wild mosquitoes were collected from up to 15 houses per village using CDC light traps [40]. The collection sites were purposively selected based one previous surveys which indicated high abundance of *An. funestus* [41]. The mosquitoes were first classified by sex and species group using morphological keys [42]. The females of *An. funestus s.l*. were then gently aspirated using mouth aspirators into 30 × 30 cm holding cages and given a 10% glucose solution at the mosquito biology laboratory at Ifakara Health Institute (i.e., the VectorSphere) for blood-feeding and rearing. This process was repeated by adding new females every week for over 26 months to support the colonization efforts and experimentation as described below. The FUMOZ colony at Ifakara Health Institute was established in 2018 using eggs supplied from the VCRL-NICD laboratories in South Africa, where it was established in 2000 and maintained for over two decades [13]. For this colony, the ‘filial generation’ therefore indicates the number of generations passed in the VectorSphere since establishment from the VCRL-NICD.

### Colonization of *Anopheles funestus* from Tanzania (FUTAZ)

Field-collected *An. funestus s.l.* females were allowed to acclimate for one day in the insectary, then all the dead adults were discarded and the remaining females were blood-fed using avian (chicken) blood [11]. This blood meal was provided to all field-collected mosquitoes only once, regardless of whether they had previously blood-fed or not. The females were kept individually in separate cups covered with netting and provided with filter papers for egg-laying (oviposition) as described by Ngowo *et al*., [11]. After the egg-laying, the species identity of the adults was re-done based using morphological keys [42] and confirmed by PCR [36], or by egg-morphology followed by PCR on a small subset of the females, to select progeny of *An. funestus s.s.* The offspring of these wild-collected females became the first generation (F1) laboratory population for the FUTAZ colony. The F1 progeny obtained were pooled into the 30 × 30cm cages for onward colonization. This process was repeated weekly, each time bringing in between 200 and 1,200 females per week, until there was stable progression past F2. After tens of several attempts, the first stable FUTAZ colony was obtained in October 2019, which required no further input of eggs from wild females. Subsequently, using these procedures, more than 30 generations of FUTAZ colony have been maintained.

The rearing of the FUTAZ and FUMOZ colonies followed the same procedures as described by Ngowo *et al*., [11], except that for FUTAZ, there were several repeated attempts before a laboratory-stable line could be selected. In brief, blood-fed adults were maintained in cages for four days, after which oviposition bowls were placed in each holding cages overnight. The eggs from the self-sustaining FUTAZ and FUMOZ lines were collected using white rectangular oviposition bowls (5 × 18.5 × 12.5 cm) containing 200 ml of tap water. Hatched larvae were fed using a finely ground mixture of dog biscuit and brewer’s yeast (ratio: 3:1) as described earlier [11]. Once the larvae had started maturing, the rearing basins were checked daily for the emerging pupae, which were collected and added into adult cages (30 × 30 cm) and maintained on 10% sugar. The first blood meal was provided 4-6 days after emergence and the blood-feeding repeated every five days there-after, *ad libitum*.

### Assessment of key fitness traits

Between 100 and 150 randomly selected adult females from each strain at different generations were used to assess mating success 15 days post emergence, body size and fecundity (Table 1). Females from the field-collected founder population (F0) were also examined regularly to keep track of the fitness traits in wild populations and compare those with the fitness of laboratory-reared mosquitoes. Data from the long-established FUMOZ colonies and also from an existing *An. arabiensis* colony (established in 2010) were used for additional comparison. The *An. arabiensis* colony was maintained using the same rearing and insectary conditions as FUTAZ and FUMOZ, the only difference being the use of TetraMin® fish food (Tetra GmbH, Herrenteich Melle, Germany) as a food source for the *An. arabiensis* larvae. The mosquitoes analyzed were obtained from different filial generations (FUTAZ: selected between F0 and F21; and FUMOZ: between F8 and F34 of maintenance since brought to the VectorSphere). These were not necessarily sampled consecutively since it was not possible to obtain adequate test material at every generation without depleting the colonies.

**Table 1:** Filial generations at which different phenotypic and genetic attributes were examined for the different mosquito colonies, populations*.

| Mosquito populations, colonies, and filial generations |  |  |  |  |  |
| --- | --- | --- | --- | --- | --- |
| Attributes tested | Wild *An. funestus* | FUTAZ | FUMUZ-Ifakara | FUMUZ-VCRL-NICD** | *An. arabiensis* |
| Percentage mated | N/A | F0 to F21 | F9 to F34 | - | Unknown |
| Wing length | N/A | F0 to F21 | F9 to F34 | - | Unknown |
| Fecundity | N/A | F0 & F6 | F16 | - | Unknown |
| Time to 50% mated | - | F9 | F16 | - | Unknown |
| Insecticide resistance | N/A | F8 | F17 | - | - |
| RFLP-type | N/A | F24 | F36 | Unknown | - |
| Clades Type | N/A | F24 | F36 | Unknown | - |

\*N/A means that the filial generations were not applicable for the wild caught *An. funestus*.
\*\*For the FUMUZ colony maintained at Ifakara, the ‘filial generation’ indicates the number of generations passed inside the VectorSphere facility since establishment from the VCRL-NICD laboratories

Mating success was measured in terms of the insemination status of females, by dissecting and microscopically observing their spermatheca [43,44]. From 1-15 days post emergence, about 20 female mosquitoes were randomly aspirated daily and dissected (Table 1). Additionally, the wing lengths of female mosquitoes were measured as a proxy for body size using calibrated micrometre ruler [45]. For each mosquito strain, to increase the chance of female mating, about 360 females and 500 male pupae (sexing based on their terminalia differences) were placed in a standard mosquito cages, and the measurements taken one day after emergence [46]. Lastly, to assess fecundity, about 150 females (per strain and generation) were provided with a blood meal via a volunteer arm, then moved into individual holding cups for three days before being transferred to oviposition cups for egg laying. The number of eggs laid by each female was recorded, excluding all females that did not lay eggs. Fecundity of wild *An. funestus* was assessed by collecting blood-fed females, holding them for three days and then providing each individual female with the egg-laying cups. In contrast to assessments of mating and body size which were done across multiple generations, fecundity data was collected at only one generation for each of the two *An. funestus* colony lines of interest (FUTAZ: F6; FUMOZ: F16).

### Assessment of insecticide resistance status

To assess whether insecticide resistance levels had changed during colonization, standard World Health Organization (WHO) test procedures for insecticide susceptibility monitoring were used [47]. The assays were done in the founder population (field collected *An. funestus* from Tanzania and in one generation of FUTAZ (Table 1). Comparative tests were performed on FUMOZ, which had originally been derived from a pyrethroid-resistant population [13].

Standard concentrations of five different insecticides from four different classes were used: pyrethroids (permethrin, 0.75% & deltamethrin, 0.05%), organochlorine (DDT, 4%), carbamate (bendiocarb, 0.1%) and organophosphate (pirimiphos-methyl, 0.25%). Resistance or susceptibility was measured based on percentage of dead mosquitoes within 24 hours after exposure. Test results were interpreted according to the WHO guidelines [47]; the mosquito population was considered susceptible if mortality exceeded 98%, possibly resistant if mortality ranged between 90% and 98%, or fully resistant if mortality was below 90%.

### Molecular characterization

Approximately 200 of randomly selected adult female mosquitoes from each colony, from different filial generations were used for molecular characterization (Table 1). For the wild FUTAZ, mosquitoes were first classified by sex and species group using morphological keys [42], and females of *An. funestus s.l.* were further identified to species level by PCR [36]. DNA was extracted from the legs or wings of female *An. funestus* and analysed by PCR to confirm species. The PCR reaction was completed inside a 0.2ml microcentrifuge tube containing 1X PCR reaction buffer (100mM Tris-HCl pH 8.3, 500mM KCl), 0.23mM dNTPs Mix, 1.38mM MgCl_2_, 0.24µM FUN, LEES, RIV, VAN, PAR and UV primers, 0.02U of *Thermus aquaticus* (*Taq*) DNA polymerase enzyme (5U/µl) and 1µl DNA template [36]. The PCR process included a denaturation step at 94°C for 2 minutes, followed by 30 cycles at 94°C for 30 seconds, annealing step at 45°C for 30 seconds, and elongation at 72°C for 40 seconds, and a final elongation of 72°C for 5 minutes. Products were visualized after electrophoresis on an ethidium bromide-stained 2.5% agarose gel. Standard positive controls were included by amplifying the DNA of known laboratory strains of *An. funestus*, and positively identified field specimens of *An*. *rivulorum* and *An. parensis*. The negative control consisted of the PCR mix without DNA template.

#### Assessing the intragenomic variations in the An. funestus s.s. populations

Digestion of the third domain (D3) of the ribosomal DNA, in combination with analysis of the internal transcribed spacer 2 (ITS2) region, have previously been used to resolve genetic differences between multiple species in both the Afro-tropical *An. funestus* group and the Asian *An. minimus* group, the two highly related but geographically separate mosquito groups [48]. The D3 region has also enabled an understanding of intragenomic variations within the *An. funestus* populations, and to study the biogeography of the species [49,50]. Here, the D3 region was amplified, and PCR used to analyse its restriction fragment length polymorphisms (PCR-RFLP), following procedures by Garros et al (2004) [48]. After PCR amplification, 10μl of the amplified D3 product was restricted and digested by using *Hpa*II enzymes. Restricted products were electrophoresed on a 3% agarose-gel in 1X TBE buffer with a PH of 8.0. For each PCR reaction, two negative controls were included, the first utilizing the "no-DNA" extraction as template and the second containing an aliquot of the PCR mixture without the template. The resulting D3 copies were used to classify samples in the FUTAZ, FUMOZ and wild *An. funestus* populations as either M, W, MW, Y or Z-types [34].

#### Analysis of the mitochondrial DNA clades

Clade analysis was used to compare the DNA sequences of the mitochondria from different strains of *An. funestus* (FUTAZ, FUMOZ and wild *An. funestus* mosquitoes) in order to understand the evolutionary relationships between them. The hydrolysis probe analysis protocol described by Choi *et al*., [37] was used, with specific probe sequences to identify *An. funestus* clades I and II. A 20μl reaction mixture was prepared containing 1X IQ-Supermix, forward and reverse primers, clade I and clade II probes, and DNA template (40 to 600ng). DNA from known clade I and clade II specimens were used as positive controls, and samples without DNA were used as negative controls. Thermocycling was conducted at 95°C for 10 minutes, followed by 45 cycles of 95°C for 10 seconds, and an extension at 63°C for 45 seconds, using a c1000tm thermal cycler. A 6-FAM probe was used to identify clade I, and a VIC probe was used to identify clade II (*Supplementary Figure-1*). Results were accepted only if the amplification curves were significantly higher than the baseline curve, all positive controls were amplified, and there was no amplification in the negative controls.

### Statistical analysis

Statistical analysis was done using R-software Version 4.0.3 [51]; to test for differences in fitness parameters of interest (mating, fecundity, body size and insecticide resistance) between different colonies (FUTAZ, FUMOZ, wild FUTAZ and *An. arabiensis*), and across successive generations within the same colony (Table 1). Generalised linear models (GLMs) were used to estimate mean values of the outcome variables (% mating, body size, fecundity) and test how they varied between colonies and filial generations. Generalized Linear Mixed Models (GLMM), using the R package, “*lme4”* [52] were used to predict time (days) needed to reach 50% mating in the cages; and to analyse variation in 24hr mortality against the five candidate insecticides. To test for directional changes in traits through time, filial generation was modelled as a continuous variable, each time fitting its effects as both linear and quadratic to also identify any non-linear effects. The function, *“dplyr”* [53] was used to subset the data for each colony so as to model them as independent data sets and avoid misleading predictions.

The binary outcome of insemination status was modelled following a binomial distribution, separately for each mosquito strain, keeping the fixed effects of interest as filial generation, the quadratic term of filial generation and wing length. On the other hand, mosquito body size data (wing length) were modelled following the gamma variate as it is classified as non-negative and positive-skewed variables. Variations in body size were modelled separately for each strain; with generation and quadratic term of generations being fixed factors. Lastly the analysis of fecundity focussed only on testing for differences between the colonies at a single time points (F6 for FUTAZ and FUMOZ after 16 generations at the VectorSphere (total generations under colonization unknown). Comparison was made between the fecundity of these strains and wild FUTAZ and *An. arabiensis* mosquitoes. Fecundity data were modelled following a negative-binomial distribution to account for the over-dispersion of the count data, including colony ID and wing length as fixed factors.

An additional analysis was performed to test for associations between the number of days spent in a laboratory cage (1-15) and the insemination status of adult females (yes or no). This was done by estimating time taken for at least 50% of females in a cage to mate. The outcome variable was insemination status (0/1) with the explanatory variables being colony ID. These measures were also taken at a single time point corresponding to F9 for FUTAZ, F16 for FUMOZ and unknown generation for *An. arabiensis*. Fixed effects of interest were colony ID and number of days post emergence, with a random effect being cage replicate, using a hierarchical/nested design.

Insecticide resistance was analysed by testing how the proportions of mosquitoes dying within 24 hours following insecticide exposure varied between different *An. funestus* strains (FUTAZ and FUMOZ) and insecticide types. These mortality data were modelled using a binomial distribution, with the outcome variable being percentage mortality and the fixed effects being colony, insecticide type, and their interaction. Experimental replicate was included as a random effect. Model selection was carried out by progressively deleting terms from the maximal model with the "*drop1()*" function, and likelihood ratio tests (LRTs) were employed to assess the relevance of explanatory variables. The “*ggeffects*” package was used to produce fitted values and 95% confidence intervals for all statistically significant terms in the best model [54]. The “*ggplot2*”, R programmes were used to create all the graphics [55].

Finally, the data on mitochondrial clades was analysed by calculating the percentage proportions of the clades detected by the specific probes in the colonies and wild populations. The same approach was used for the data from PCR-RFLP analysis of the D3 region.

## Results

### Mating success

In all colonies, the ‘generations’ had a curvilinear relationship with mating success, measured as proportion of inseminated females (Figure 1). There was an initial dip for the first few filial generations followed a gradual rise. The percentage insemination in FUTAZ decreased from 68.2% at F0 to 53.4% at F6, before consistently increasing to 92.1% by F21 when the monitoring ended. In the long-established FUMOZ colony, the percentage of insemination remained high and stable for the first 20 generations (73-84%) and slightly increased to 91.5% during the final period of monitoring (Figure 1).

**Figure 1:**
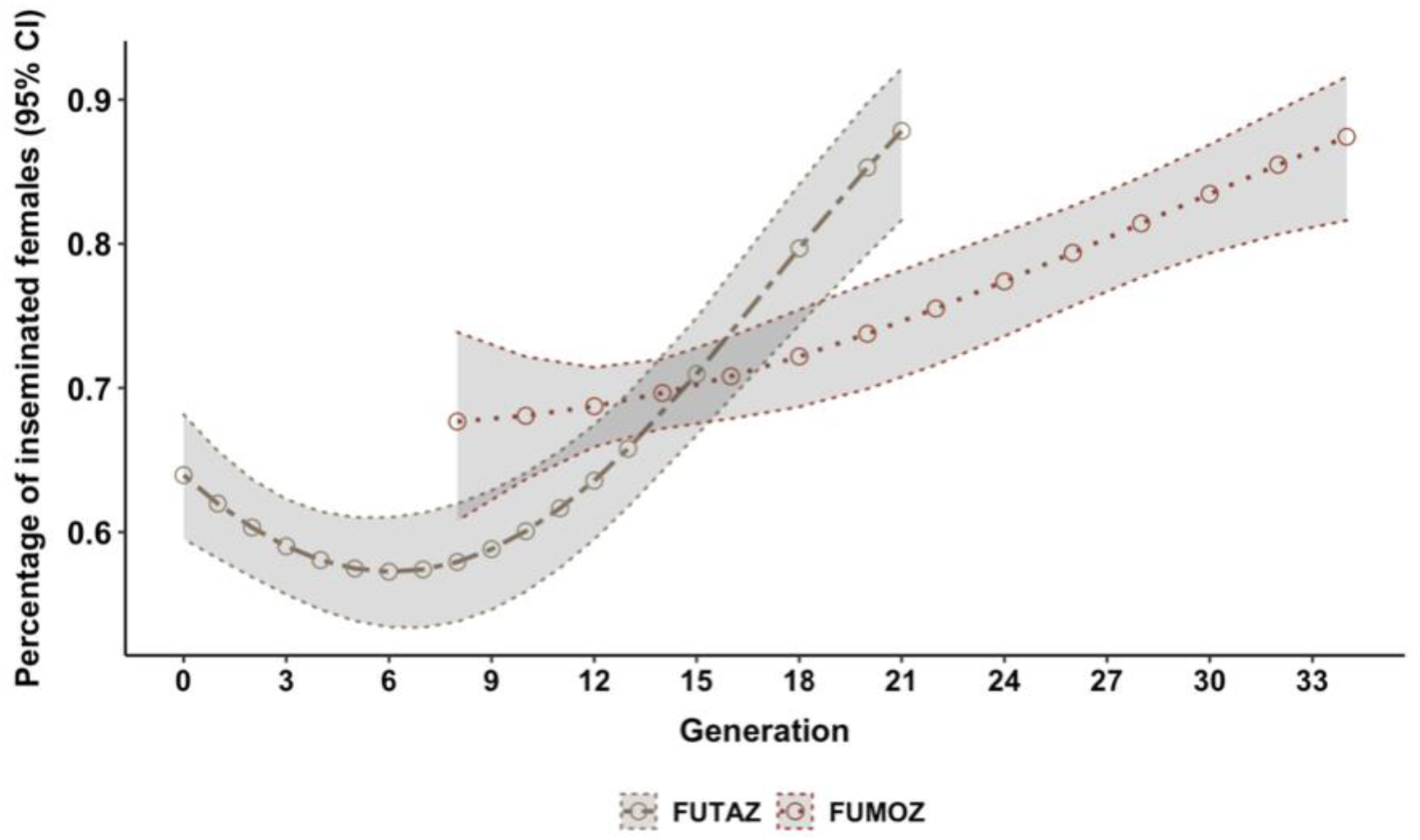
Mean predicted probability of inseminations in three different *Anopheles funestus* colonies at different generations of colonization. “Generation’ indicates the number of generations passed in the Ifakara Health Institute mosquito biology laboratory, (the VectorSphere) since colony establishment. For FUTAZ, F0 was the field-collected wild mosquitoes, while for FUMOZ, it refers to mosquitoes obtained from the VCRL-NICD laboratory, where the colony was already long-established. The shaded regions around the predicted line of each colony represents the 95% confidence interval (CI).

Within a given generation, the probability of insemination increased with the number of days females spent in laboratory cages post emergence in each of the *Anopheles* colonies, (Colony/Day (χ^2^ =823.22, df=5, *p*<0.001) (Figure 2). Insemination was fastest in the *An. arabiensis* colony, in which 50% of females were inseminated within 4.8 (95%CI: 4.0-5.3) days post emergence. In contrast, it took more than 10 days to reach 50% insemination in both the long-established FUMOZ [11.7 days (95%CI: 10.8-12.5)] and the newly established FUTAZ colony [13.1 days (95%CI: 12.3-14.2)] (Figure 2).

**Figure 2:**
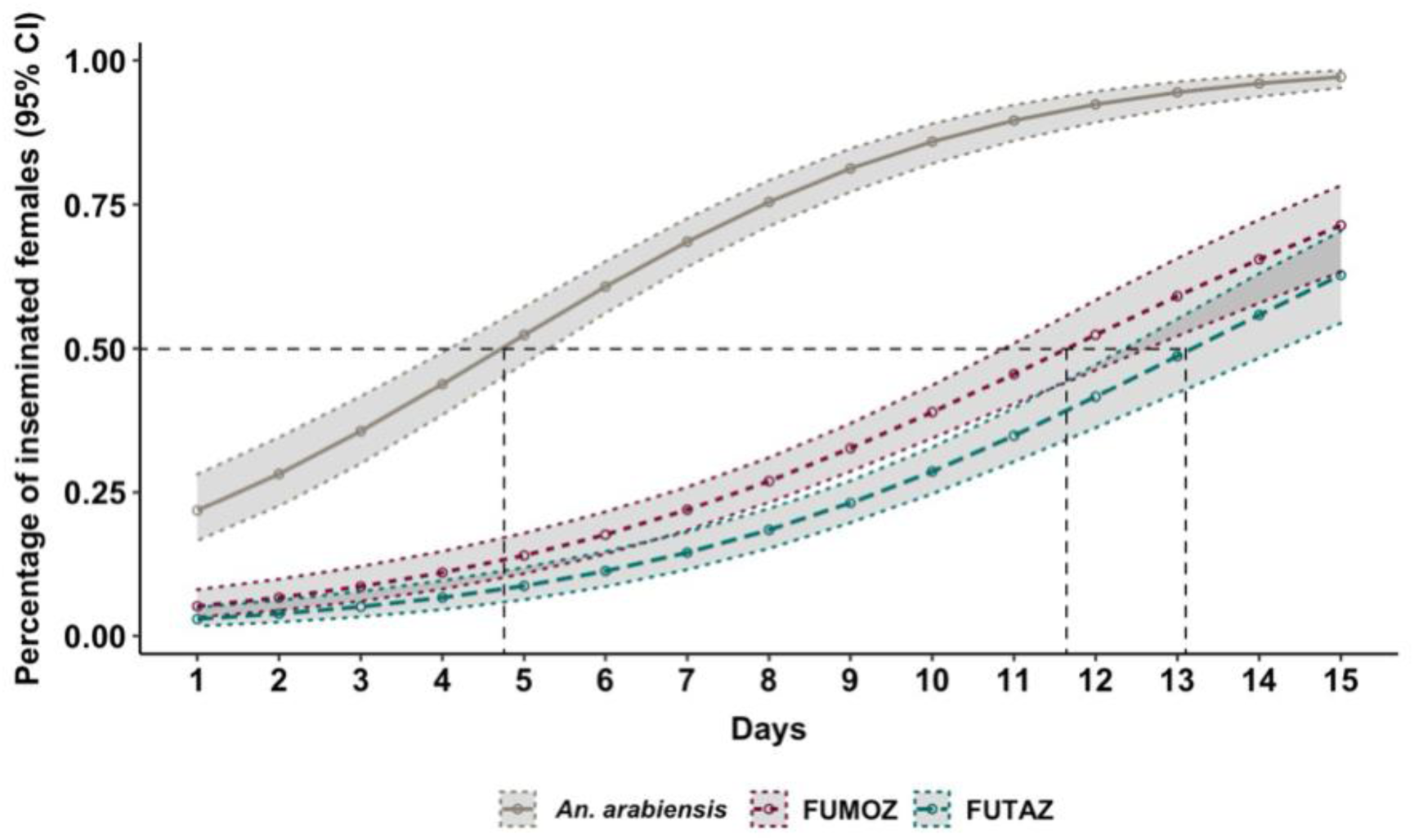
Mean predicted insemination rate of female *Anopheles* mosquitoes in the different colonies (FUTAZ, FUMOZ and *An. arabiensis*). Dotted lines represent the number of days needed to achieve 50% mating success in the cage for each strain. The shaded regions around the predicted line of each colony represents the 95% confidence interval (CI).

### Body size

The body size of *An. funestus* (measured from wing lengths as a proxy) also had a curvilinear association with generation in the FUTAZ colony, though this relationship was linear in the FUMOZ colony (Figure 3). In FUTAZ, wing length decrease from approximately 2.8mm at F0 to 2.6mm at F12, before increasing again to reach 2.7mm by F21. In FUMOZ, the wing lengths increased consistently with generations from 2.6mm at the start to 2.8mm at the end of monitoring period (Figure 3).

**Figure 3:**
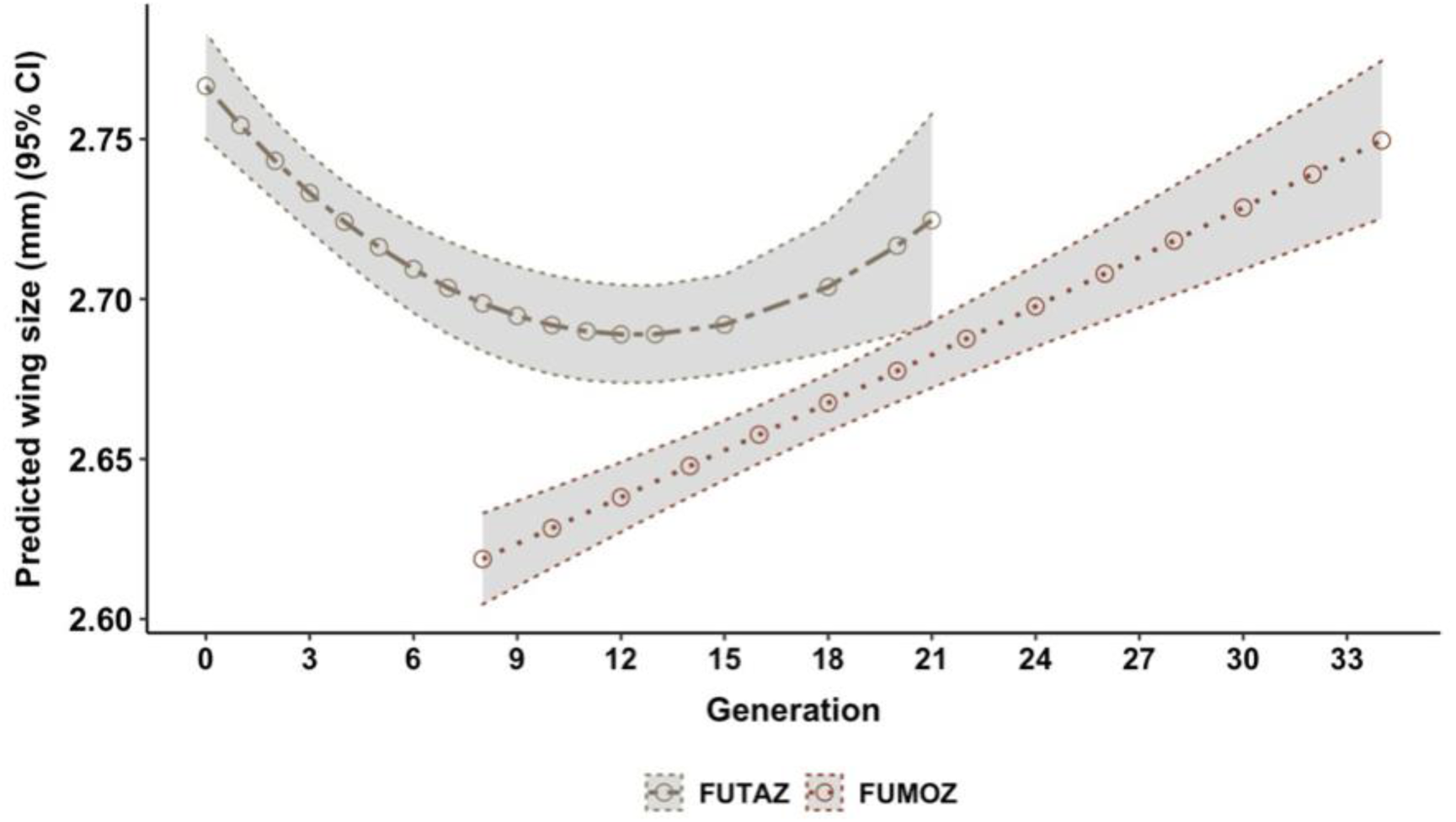
Mean predicted wing length (mm) in different *Anopheles funestus* colonies at different generations of colonization in the VectorSphere.

### Fecundity

Generally, the wild *An. funestus* laid an average of 98 (95% CI: 91-106) eggs, while FUTAZ mosquitoes laid 74 (95% CI: 63 - 92) and FUMOZ females laid 80 (95% CI: 73 - 90) eggs per female (Figure 4a). The number of eggs laid by *An. arabiensis* was 76 (95%CI: 63-92). The primary comparison of interest was how fecundity varied between the relatively “new” FUTAZ colony versus the “older” *Anopheles* colonies (FUMOZ and *An. arabiensis*), rather than how it varied across generations within the same colony. In this analysis, fecundity was best described in a final model that included the interaction between wing-length and colony-line (wing-length: colony-line; χ^2^ =55.73, df=7, *p*<0.001). In all colonies, fecundity was positively associated with body size (wing length) as bigger mosquitoes tended to lay more eggs than smaller mosquitoes (Figure 4b). The strength of this relationship was however varied by colony. Generally, the laboratory-reared *An. funestus* laid 61 to 99 eggs, with no significant difference between colonies (Figure 4).

**Figure 4:**
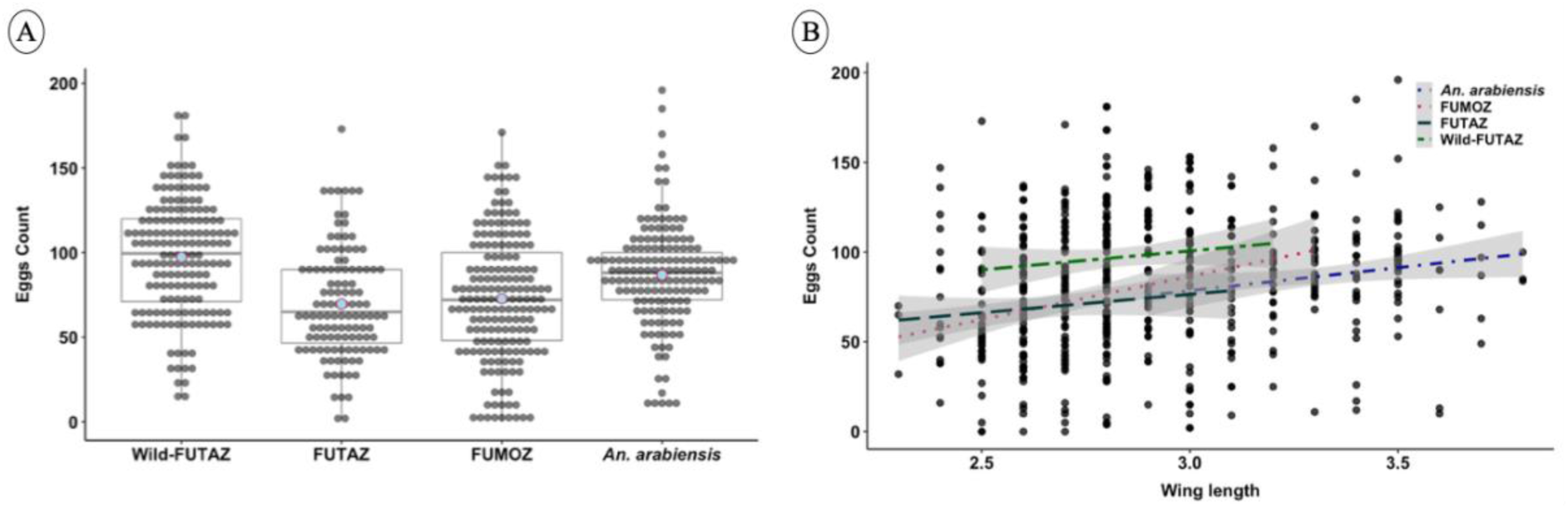
Fecundity of female mosquitoes: (a) predicted mean number of eggs laid by wild *Anopheles funestus*, FUTAZ (F6), FUMOZ (F16) and *Anopheles arabiensis* in the laboratory, and (b) relationship between the body size (wing length) and the number of eggs laid by mosquitoes from the different laboratory colonies

### Insecticide resistance

When the mosquitoes were exposed to permethrin, the post-exposure mortality was generally lower in the wild founder population of *An. funestus* (26%) and the new FUTAZ colony (43%), than in the long-established FUMOZ colony (89%). A similar yet less extreme trend was observed for DDT, where post exposure mortality was 84% for the field-collected mosquitoes, 99% for FUTAZ colony and 100% in FUMOZ females (Figure 5). However, there was no consistent trend following exposure to deltamethrin, against which resistance was generally similar across colonies (Figure 5). High post-exposure mortalities were observed in all colonies following exposure to bendiocarb and pirimiphos-methyl, indicating all the *An. funestus* populations were susceptible to these insecticides (Figure 5).

**Figure 5:**
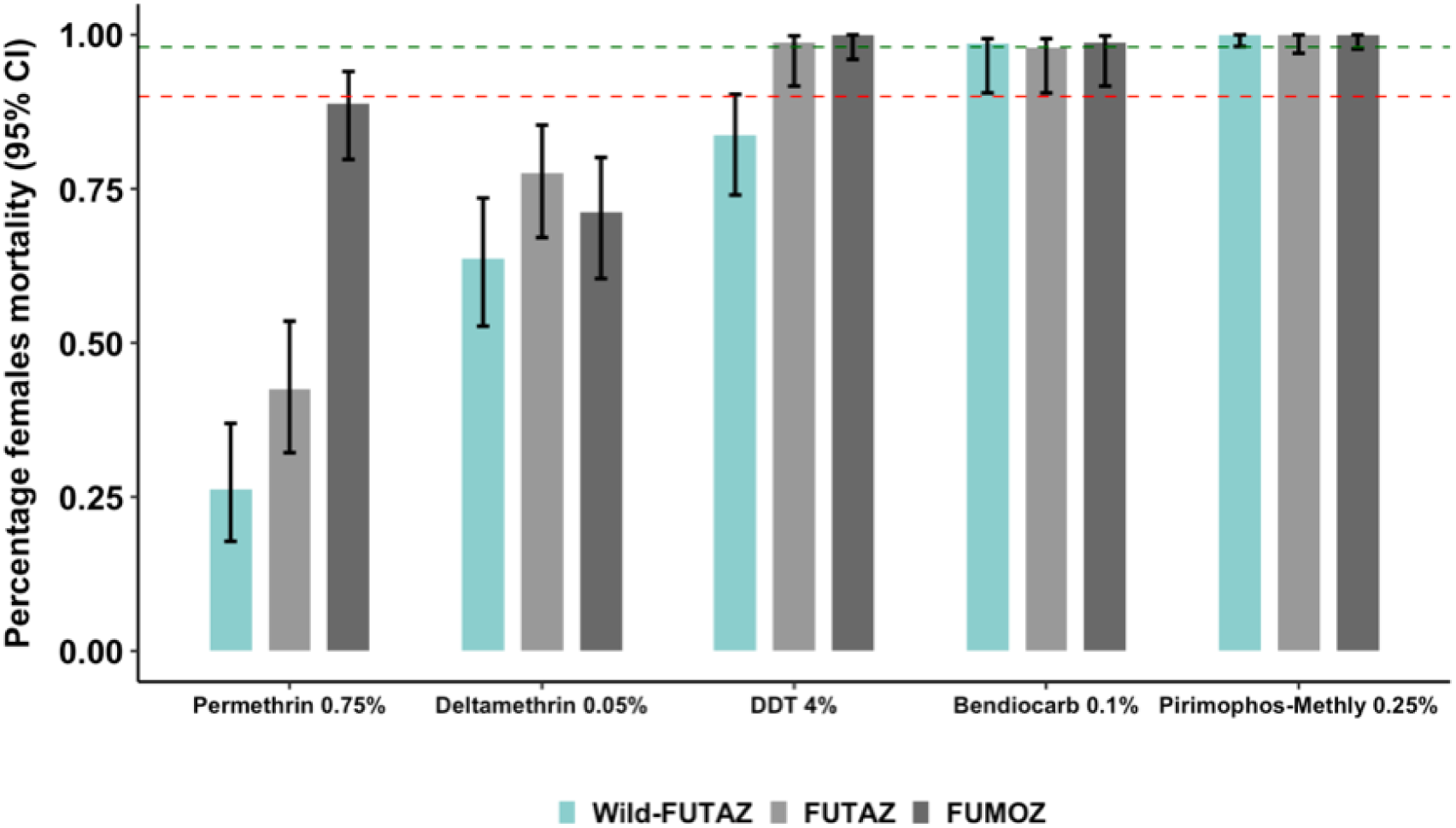
Mean predicted mortality (and 95% confidence intervals (CI)) of different *Anopheles funestus* colonies and filial generations in standard WHO susceptibility tests. The dotted lines from 90% (red dotted line) to 98% (blue dotted line) indicate thresholds for possible resistance and complete resistance respectively.

### Mitochondrial clades analysis

All the specimen from different colonies were re-confirmed by PCR as being *An. funestus* before the clades assay. Additionally, the TaqMan^®^ assay used to identify the mitochondrial clades showed that 89.4% of the wild-collected *An. funestus* samples from Tanzania belonged to Clade I (Table 2), while only 7.6% were of clade II. On the other hand, the FUMOZ mosquitoes were mostly of Clade II (88.5% in the NICD-VCRL laboratories and 64.5% in the Ifakara Health Institute Laboratory). The FUTAZ colony was 80.4% Clade I, thus being more closely related to the wild Tanzanian *An. funestus* than FUMOZ (Table 2).

**Table-2:**
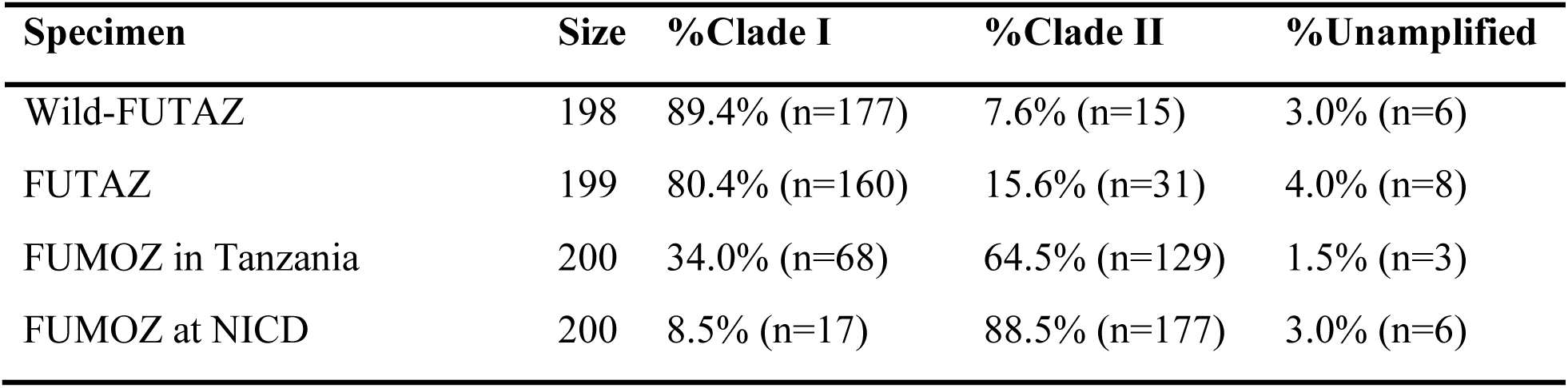
Clade composition of the different *An. funestus* populations and colonies.

### Intragenomic variations based on the PCR-RFLP assay of the D3 domain

Analysis of the intragenomic variation of the D3 gene, using the *Hpa*II RFLP-PCR, revealed that more than 90% of the field-collected *An. funestus* mosquitoes from Tanzania were “Y” type, while FUMOZ mosquitoes were mostly “MW” type (Table 3). In the FUMOZ colony maintained in Tanzania, 91.5% were “MW” Type and only 3% were “Y” type. Similarly, in the FUMOZ colony maintained at the VCRL-NICD in South Africa, 90.5% were “MW” Type and only 0.5% were “Y” type. The FUTAZ colony (90.5% “Y” Type) was therefore more closely related to the wild-collected *An. funestus* (91.4% “Y” type) than to the FUMOZ colonies, which were mostly “MW” types (Table 3; *Supplementary Figure-2*).

**Table-3:** Summary table of the intragenomic variation of the D3 ribosomal DNA gene of An. funestus using the PCR-RFLP.

| Specimen | %Y-type | %M-type | %MW-type | %Unamplified |
| --- | --- | --- | --- | --- |
| Wild-FUTAZ | 91.4% (n=180) | 4.0% (n=8) | 0.0% (n=0) | 4.6% (n=9) |
| FUTAZ | 90.5% (n=180) | 4.0% (n=8) | 0.0% (n=0) | 5.6% (n=11) |
| FUMOZ in Tanzania | 3.0% (n=6) | 0.0 (n=0) | 91.5% (n=183) | 5.5% (n=11) |
| FUMOZ at NICD | 0.5% (n=1) | 0.0% (n=0) | 90.5% (n=181) | 9.0% (n=18) |

## Discussion

Difficulties in the laboratory colonization have greatly hindered scientific studies on the basic biology and ecology of important malaria vector species, notably *An. funestus*. Relative to other vector species, the resulting lack of a comprehensive understanding of *An. funestus* has delayed the development of effective control measures. Fortunately, after numerous attempts, we were able to successfully colonize *An. funestus* from wild in rural Tanzania. The new colony of *An. funestus* from Tanzania is named FUTAZ, as opposed to the long-established colony of *An. funestus* from Mozambique, i.e., FUMOZ. This paper is the first description of how this species adapted to laboratory colonization over multiple generations.

Mosquito colonization challenges have previously been attributed to the poor reproductive success of the F1 generations [9,11]. In some eurygamic species, laboratory colonies were successfully established through forced mating of the first few generations, which subsequently adapted and became more stenogamic (i.e. capable of free mating in restricted environment) [17]. This current approach did not involve forced mating, but instead repeated collection of large numbers of wild females (founder population ‘F0’) and obtained their offspring ‘F1’ using the procedure of individual oviposition in small paper cups to select *An. funestus* [11]. The repeated addition of wild-caught females into the founder population cages was stopped only after the colony was judged stable on the basis of the F2 females laying sufficient eggs for further generations (after two years of trials). Due to the need for frequent field collections, close proximity to rural villages with high densities of *An. funestus* was a crucial element in the success of the FUTAZ colony.

Earlier attempts to colonize *An. funestus* in Tanzania were impeded by multiple bottlenecks in several crucial life cycle processes including reduced mating, reduced fecundity and slow larval development [11]. In this current study, insemination rates of *An. funestus* were also considerably reduced within the first few laboratory generations (F1-F6), matching observations reduced mating in F1 and F2 *An. funestus* in Tanzania [11], and in other eurygamic species, including *An. marajoara* in Brazil [17]. The timescale over which *An. funestus* began to show signs of adaptation to stenogamy here is also consistent with studies of other eurygamic species. For example, it took six generations before *An. albitarsis* from Brazil transitioned to free mating in laboratory colonies (F6), leading to their successful colonization for more than 18 generations [56]. Similar observations were reported for *An. maculatus*, where, after forced mating in the initial stages, the colony became well adapted and reached more than 85% mating success after five generations [57]. Given the mating-related bottlenecks, it is important to ensure that the initial founder population of a laboratory colony is large enough to persist through this early drop in mating success (F1 to F7). In this current study, the newly colonized FUTAZ strains exceeded 75% mating success after completing the first 15 generations in the laboratory. This matched the performance of the long-established FUMOZ colony, suggesting that FUTAZ had crossed the stability threshold.

In field settings, majority of African malaria vector species are known to mate within the first few days after emergence, usually before their first blood meal [58]. Mating is however likely to take longer in laboratory cages for eurygamic species like *An. funestus*. This is an important determinant of colony stability, as the longer it takes for mosquitoes to mate, the more likely they will die before producing eggs for the next generation. This current study found that mating occurs very slowly in laboratory cages of *An. funestus*. It took more than ten days post-emergence for 50% of *An. funestus* females to be mated, compared to just five days in *An. arabiensis*. The reason for the delayed mating in *An. funestus* remains unclear but may reflect their more eurygamic behaviour in general, though in early filial generations it also reflects a general lack of acclimatization to the laboratory environments. These findings therefore suggest that although the mosquitoes eventually adapt to in-cage mating, females do this only at much older ages.

Mosquito body size is also commonly associated with their fitness [11,59]. For example, mosquito wing length has been positively associated with adult survival [60–62], mating success [63], fecundity [11] and energetic reserves [64]. This study found that mean wing lengths of *An. funestus* during the early generations of colonization were lower than in wild populations. It is often assumed that laboratory mosquitoes are larger than field mosquitoes, because of the greater stability and supply of nutrition in laboratory studies. The reduction in *An. funestus* body size observed during the first few laboratory generations may also have been responsible for the lower mating success and fecundity of the F1-F6 generations. Colonization has the potential to change the body sizes of mosquitoes through adaptation to the larval nutrition [59], or indirect selection that bigger mosquito have higher mating likelihood in the laboratory [24]. In this study, mosquito body sizes for FUTAZ dropped consistently for the first seven generations (F1-F7), and started to increase from F8 to end at about same size of FUMOZ colony. Eventually, the mosquitoes kept in the laboratory colonies achieved mean body sizes that were either equal to or lower than the sizes of the wild-caught founder females.

Another barrier to laboratory colonization of *An. funestus* is poor egg production in captivity. Here the mean fecundity of the new and old *An. funestus* colonies were examined after at least six generations (F6) inside the laboratories. There was no statistical difference in the number of eggs produced by wild-caught *An. funestus*, FUTAZ, FUMOZ and the *An. arabiensis* colony. Although we cannot be certain egg production was not reduced during the early generations of colonization, these results indicate that fecundity may not be limiting beyond the first few generations in colony. The mean fecundity of Tanzanian *An. funestus* observed here (wild *An. funestus*: 97.2 ± 33.9 eggs/female; FUTAZ: 69.6 ± 32.1 eggs/female) are similar to that reported for the FANG strain, another long-established *An. funestus* colony (67.1 ± 17.1 eggs/female) [12]. Even after extensive laboratory adaptation, the number of eggs laid by *Anopheles* can often be fewer than by wild mosquitoes [12].

This study also investigated the impact of colonization of insecticide resistance. The founder populations from which the FUTAZ colony was derived is known to be highly resistant to pyrethroid insecticides [65]; mirroring the high pyrethroid resistance of *An. funestus* in many other places [13,20,66]. The FUMOZ, which originated in Mozambique, was also known to have high levels of pyrethroid resistance at VCRL-NICD [13,67]. Despite the general hypothesis that resistance would decline during colonization, resulting in higher post-exposure mortalities of colonised compared to wild type mosquitoes, previous studies of the FUMOZ mosquitoes have shown that pyrethroid resistance pressure may not always lead to a regression of resistance alleles since it does not compromise fitness [68]. In this current study, the expectation that the resistance phenotype would be strongest in the wild *An*. *funestus* followed by newer colonies then long-established *An. funestus* colonies was only partially met. The new FUTAZ colony was still resistant to pyrethroids at the F6 generation, indicating this trait was not lost rapidly. However, based on the percentage mortalities observed, there was a trend of lower resistance with ‘time since colonization’ across the three *An. funestus* populations tested. The post-exposure mortality from permethrin was lowest in the wild founder population, intermediate in the FUTAZ line, and highest in the FUMOZ colony. The high post-exposure mortality of the FUMOZ colony exposed to permethrin indicates resistance has been lost during it’s nearly two decades of laboratory colonization. In contrast, a different pattern was observed for deltamethrin 0.05%, with almost similar post exposure mortality, which negates the general the hypothesis of declining resistance. These observations might reflect differential responses of mosquitoes to Types I and II pyrethroids. All colonies were fully susceptible to bendiocarb 0.1% and pirimiphos-methyl 0.25%, consistent with previous findings from Tanzanian field populations [65]. Molecular assays are therefore needed to clarify these differences and whether there is a continuous trend of resistance loss, as observed in other *Anopheles* species [68].

Mitochondrial DNA (mtDNA) clades analysis found that most of the FUTAZ mosquitoes belonged to Clade I, thus closely resembling the wild Tanzanian *An. funestus* populations which also belonged mostly to Clade 1 (80.4% and 89.4%). However, the FUMOZ colonies maintained in Tanzania and in South Africa were both predominantly of Clade II (64.5% and 88.5% respectively). Previously, Michael *et al*., 2005 [38] and Choi *et al*., 2013 [37] also indicated that, from Tanzania the majority *An. funestus* mitochondrial data belonged to Clade I, while those from Mozambique (where FUMOZ had originated) were clade II. The low genetic-level homoplasy (a phenomenon of convergent evolution, where different groups of organisms independently evolve similar characteristics or adaptations to similar environments or ecological niches) might suggest the bottlenecks but not necessarily rule out colonization success.

There can be considerable genomic variations within, rather than between population samples from the same or different geographic locations. It was previously observed that *An. funestus* populations can be classified into three groups by microsatellite allele frequencies: east, west, and central [33], with the eastern population including mosquitoes from the coasts of Tanzania, Mozambique, and Malawi. In sub-Saharan Africa, there are also five RFLP microsatellite types known: M, W, MW, Y, and Z [33,34]. While the Tanzanian mosquitoes, both wild and colonized, were mostly “Y” type, the FUMOZ were mostly MW type. The “Y” RFLP types had not previously been observed in south-eastern Tanzania, where the mosquitoes in this study originated. The presence of few “M” type mosquitoes in FUTAZ colony and the field-collected Tanzanian *An. funestus* populations may indicate the presence of a population structure within the area where majority are “Y” RFLP type. The same may be true for the observed presence of few “Y” type specimen among the FUMOZ colony mosquitoes. This also suggests that RFLP analysis may not be sufficiently sensitive to detect autosomal DNA variation [34].

Even though this study addressed all initial objectives, there were several limitations that might impact interpretation and generalizability. First, the fitness features associated with reproduction were only evaluated in female, not males. Male mosquitoes play a vital part in reproduction since they must be capable of surviving long enough to mate and competing successfully for females. Some of the fitness outcomes investigated here, such as female mating success, are highly dependent on male fitness. Future studies should therefore include both sexes for more comprehensive assessment of barriers to colonization. Other limitations include the absence of assessments on adult survival, larval development time and pupation, all of which may be determinants of fitness and laboratory adaptation [69].

## Conclusion

Mosquito colonies are essential for vector biology and control research because they provide a consistent and standardised source of mosquitoes for experiments. Unfortunately, *An. funestus*, a major Afro-tropical malaria vector, has historically been difficult to colonize in laboratories, thus significantly hindering research on its biology and control. This study has reported the colonization and characterization of the first laboratory *An. funestus* strain from Tanzanian (FUTAZ). The study identified the key determinants that contribute to effective colonization and the repeatability of the process. Poor insemination success and small body sizes during the early generations of laboratory maintenance were identified as major barriers to establishing stable colonies. However, by repeatedly collecting large numbers of offspring from wild females to create a larger ‘F1’ populations, it was possible to overcome these barriers and maintain the colonies through these initial generations. Within seven generations of the F1 population, there were significant improvements in both mating success and body size, which eventually matched those of a strain that had been successfully colonized for over 10 years. Molecular characterization of these mosquitoes demonstrated the genetic similarities between the FUTAZ mosquitoes and wild-caught Tanzanian *An. funestus* - as they were mostly of mitochondrial Clade 1 as opposed to Clade II and of RFLP Y-type, but also showed that the new colony can be distinguished from the FUMOZ mosquitoes, which were mostly of Clade I and RFLP type MW. As of this writing, the FUTAZ colony has now been maintained for more than 30 generations at the Ifakara Health Institute laboratories in Tanzania and will be a valuable resource for researching the biology and ecology of *An. funestus*, as well as developing effective control measures.

## Acknowledgements

We would like to thank village leaders and community members in Tulizamoyo and Sululu Villages for allowing their houses and areas to be used during mosquito collection. We would like also to express our gratitude to all colleagues in the Outdoor Mosquito Control (OMC) group at Ifakara Health Institute (IHI), who provided valuable assistance throughout this work.

## Authors’ contributions

EEH, LLK, HMF and FOO designed the study. EEH, DMM, RMN, LLM, IHN, DMM, NKN & JPM performed laboratory experiments. EEH, HMF and FOO wrote and revised the manuscript. EEH, HMF and HSN performed data analysis. MPZ, NFK, JOO, SAM, PPC, NJG, INL, SSK, DWL, BBT, EWK, PS, FT, CSW, FB, HMF, LLK & FOO reviewed the manuscript. All authors read and approved the final manuscript.

## Availability of data and materials

All data generated from this study will be available from the corresponding author as per request.

## Ethical approval

Ethical approval for this study was obtained from Ifakara Health Institute Institutional review board with certificate number (Ref. IHI/IRB/No: 26-2020) and from the Medical Research Coordinating Committee (MRCC) at the National Institute for Medical Research-NIMR (Ref: NIMR/HQ/R.8a/Vol. IX/3495). Individual verbal and written consents were also obtained from household owners whereby CDC light traps were set inside the houses for collecting adult female *Anopheles funestus* mosquitoes and verbal consent for the arm-feeders in the insectary.

## Consent for publication

This manuscript has been approved for publication by the National Institute of Medical Research, Tanzania (Ref. No: NIMR/HQ/P.12 VOL XXXV/126).

## Supplementary Materials

**Supplementary Figure 1:**
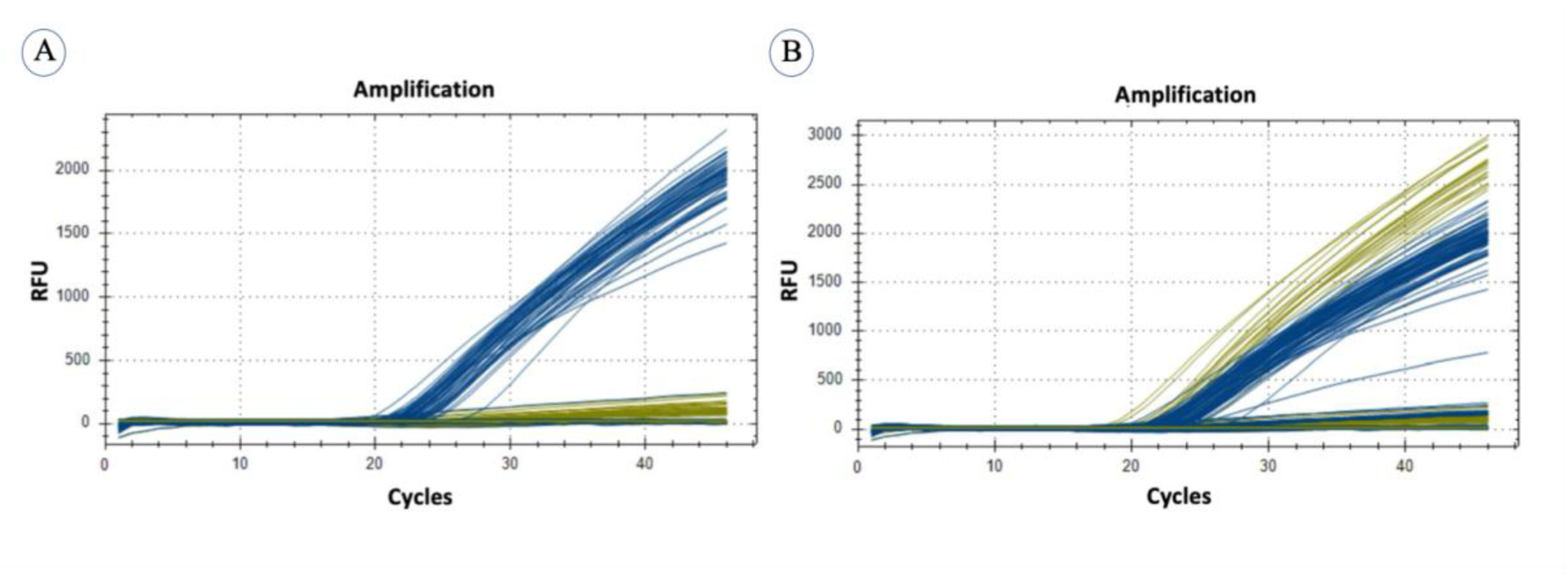
Fluorescent results from the Hydrolysis probe analysis for both FUTAZ and FUMOZ. Quantitative peaks **a)** Clade I; **b)** Clade I and II. Clade I are represented by blue peaks while Clade II indicated by yellow peaks.

**Supplementary Figure 2:**
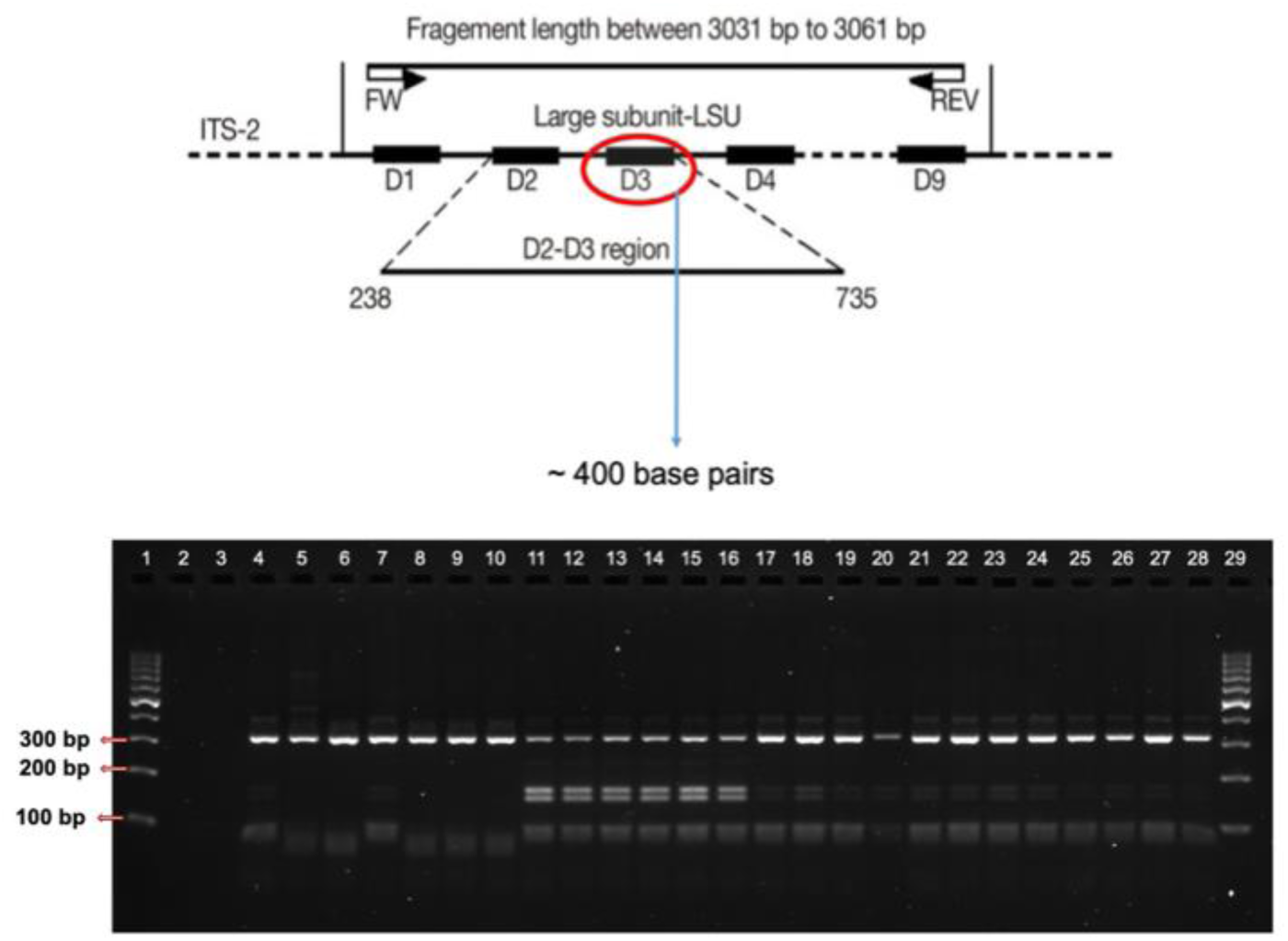
Image of the Gel electrophoresis product after digestion of the D3 region of the Ribosomal DNA; Lane 1 & 29 - Molecular marker (100-bp ladder), Lane 2 & 3 - DNA extraction & PCR negative control respectively, Lane 5-6, 8-10 - M-type, Lane 11-16 - MW-type & Lane 4,7,17-28 - Y-type.

